# Interactions between earliest *Linearbandkeramik* farmers and central European hunter gatherers at the dawn of European Neolithization

**DOI:** 10.1101/741900

**Authors:** Alexey G. Nikitin, Peter Stadler, Nadezhda Kotova, Maria Teschler-Nicola, T. Douglas Price, Jessica Hoover, Douglas J. Kennett, Iosif Lazaridis, Nadin Rohland, Mark Lipson, David Reich

**Author notes:** these authors contributed equally to this work (Alexey G. Nikitin, Peter Stadler, Nadezhda Kotova).

## Abstract

Archaeogenetic research over the last decade has demonstrated that European Neolithic farmers (ENFs) were descended primarily from Anatolian Neolithic farmers (ANFs). ENFs, including early Neolithic central European *Linearbandkeramik* (LBK) farming communities, also harbored ancestry from European Mesolithic hunter gatherers (WHGs) to varying extents, reflecting admixture between ENFs and WHGs. However, the timing and other details of this process are still imperfectly understood. In this report, we provide a bioarchaeological analysis of three individuals interred at the Brunn 2 site of the Brunn am Gebirge-Wolfholz archeological complex, one of the oldest LBK sites in central Europe. Two of the individuals had a mixture of WHG-related and ANF-related ancestry, one of them with approximately 50% of each, while the third individual had approximately all ANF-related ancestry. Stable carbon and nitrogen isotope ratios for all three individuals were within the range of variation reflecting diets of other Neolithic agrarian populations. Strontium isotope analysis revealed that the ~50% WHG-ANF individual was non-local to the Brunn 2 area. Overall, our data indicate interbreeding between incoming farmers, whose ancestors ultimately came from western Anatolia, and local HGs, starting within the first few generations of the arrival of the former in central Europe, as well as highlighting the integrative nature and composition of the early LBK communities.

## INTRODUCTION

The *Linearbandkeramik* or Linear Pottery culture (LBK) played a key role in the Neolithization of central Europe. Culturally, economically, and genetically, the LBK had its ultimate roots in western Anatolia, but it also displayed distinct features of autochthonous European Mesolithic hunter-gatherer societies. Several models for the origins of the LBK culture have been proposed over the years^1^.

The Indigenist model suggests the LBK was founded through the adaptation of elements of the West Asian Neolithic Package by indigenous Mesolithic populations exclusively through frontier contact and cultural diffusion. The Integrationist model views the formation of LBK as the integration of Mesolithic hunter-gatherers into an agro-pastoral lifeway through mechanisms such as leapfrog colonization, frontier mobility and contact. According to this model, small groups associated with the Starčevo-Körös-Criş (SKC) culture, the likely LBK predecessors in Europe, left their homelands in the Balkans (where most of their own ancestors had arrived earlier from Anatolia), and settled new areas to the northwest. Contacts with local Mesolithic groups and exchange of products would have resulted in the co-optation of hunter-gatherers into farming communities, where they would have adopted farming practices^1^. Evidence of such interactions exists at the Tiszaszőlős-Domaháza site in northeastern Hungary, containing interments of individuals of mostly hunter gatherer genetic ancestry buried in a clearly SKC context^2,3^.

The Migrationist model suggests that a sparsely populated territory of Mesolithic central Europe was taken over by pioneering agro-pastoral groups associated with the SKC culture, which gradually displaced indigenous hunting-gathering populations, who did not significantly influence the arriving Starčevo colonizers. According to this model, newcomers would have replicated their ancestral material culture in the newly settled territory without incorporating the material culture features of the local indigenous populations. Some variation, due to innovation and adaptation to the new environment and sources, would have involved changes in technology such as pottery and building material as well as lithic tool sources. At the same time, symbolic systems, such as decorative designs and cultural objects, would have remained unchanged. This model appeared at the end of the 1950s^4^ and gained wide support in the second half of 20^th^ century^5–8^.

To date, ancient DNA (aDNA) studies have convincingly shown that Neolithic European farming populations were primarily genetic descendants of central and western Anatolian Neolithic farmers (ANFs)^9–11^. Their genetic signature is clearly distinct from autochthonous Mesolithic European hunter gatherers (HGs) of central Europe (WHGs) at the level of uniparental markers such as mitochondrial DNA (mtDNA) and Y chromosome as well as genome-wide. Nevertheless, the extent to which the newcomers interacted both culturally and genetically with local hunter gatherers remains unclear; that is, it remains unclear to what extent an Integrationist or Migrationist model is accurate. Genetically, Neolithic central European farmers carried a minor proportion of genetic ancestry characteristic to WHG populations, but the extent and the timing of the WHG admixture in the gene pool of the European Neolithic descendants of Anatolian farmers varies across central Europe^3^. While the amount of WHG ancestry in European Neolithic farmers had been observed to increase throughout the Neolithic in the present-day territories of Hungary, Germany and other regions of Europe^3,9,12–15^, the initial degree of exchange remains unresolved, in part due to a scarcity of human remains contemporaneous with the earliest stages of the Neolithic farming migration.

The Brunn 2 archaeological site, part of the Brunn am Gebirge, Wolfholz archaeological complex south of Vienna, Austria (Figure 1), is the oldest Neolithic site known in Austria and one of the oldest in all of central Europe. It belongs to the earliest stage of the development of LBK, called the Formative phase. Radiocarbon dates obtained for Brunn 2 time the site to about 5670-5350 cal BCE^16–20^. The main characteristic of the settlements of the Formative phase is the absence of fine pottery and the use of coarse pottery with clear Starčevo features. The leading role of Anatolian migrants in the formation of cultural attributes of the earliest farmers of Europe is evident through the comparative typological analysis of material culture artifacts from the Brunn 2 site^20^. In addition to rich trove of culture artifacts, Brunn 2 yielded four human burials. The initial radiocarbon dating of the remains confirmed these to be contemporaneous with the earliest phase of the Brunn am Gebirge complex^20^ and, thus, to represent some of the earliest central European Neolithic farmers. We set out to perform a bioarchaeological analysis of these individuals to examine genetic ancestry as well as diet and mobility at the dawn of the European Neolithization, in an effort to refine the model of the establishment of farming in the Neolithic central Europe.

**Figure 1.**
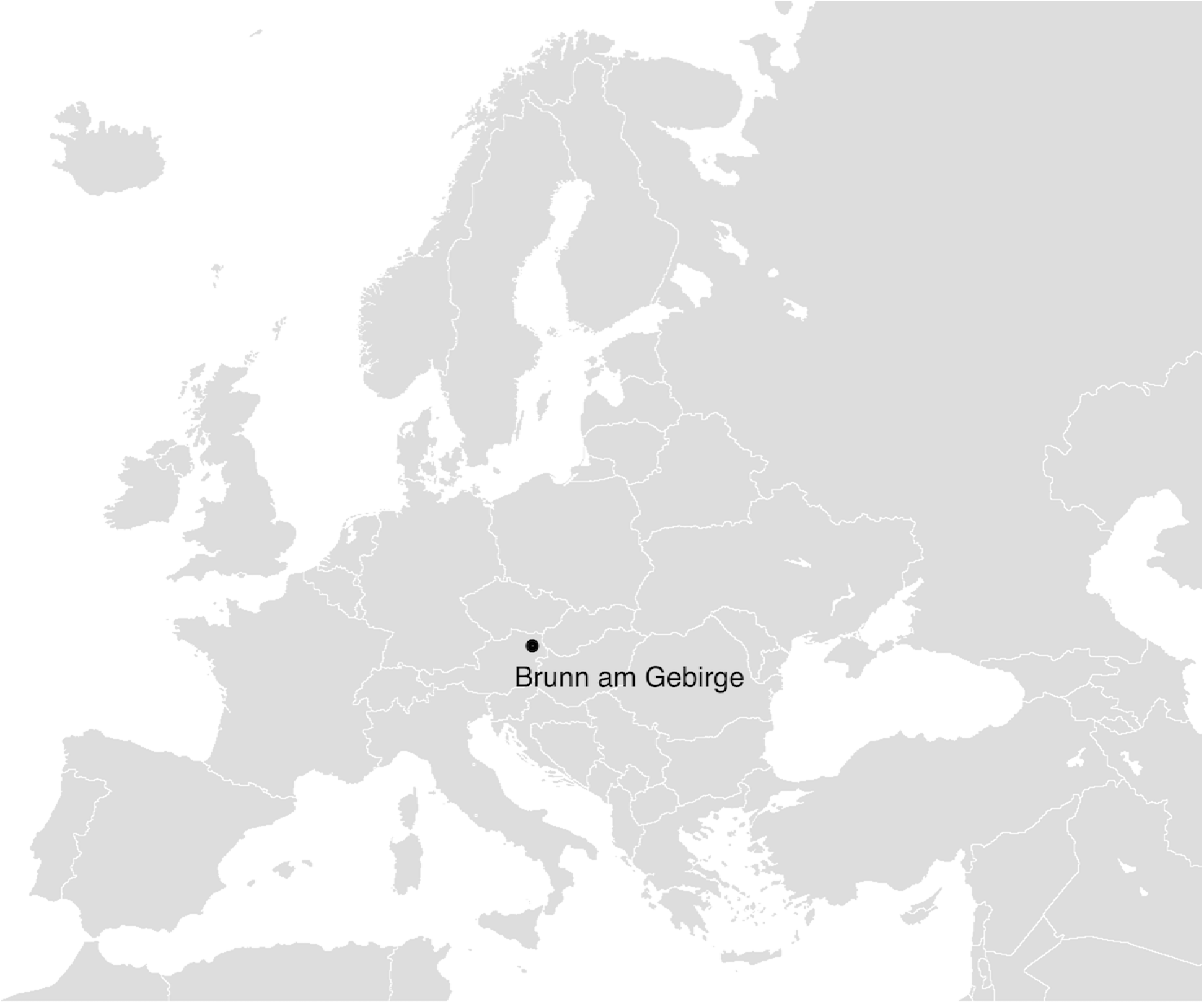
The location of Brunn am Gebirge, Wolfholz site on the map of Europe. Map image is from https://en.wikipedia.org/wiki/File:Europe_blank_map.png (image is in the Public Domain).

## MATERIALS AND METHODS

### Brunn 2 archaeological site

The Brunn am Gebirge - Wolfholz excavation site (48.120396, 16.291722) is located near Vienna, Austria. Six sites uncovered at the location have shown a development of the LBK from the Formative phase until the Musical Note Pottery (*Notenkopfkeramik*) phase. Sixteen radiocarbon dates obtained for Brunn 2 date the oldest part of this site to about 5,670-5,450 calBCE^16–20^, which places this site along with Szentgyörgyvölgy-Pityerdomb^21^ and Zalaegerszeg-Andráshida^22^ in Hungary within the Formative Phase of LBK. The main characteristics of these settlements is the absence of fine pottery and the use of only coarse pottery with clear Starčevo features. In total, 3,620 pottery vessels have been recovered from Brunn am Gebirge along with 100,000 pottery sherds. The ceramic collection from Brunn 2 includes 11 narrow-neck vessels (“amphorae”) and fragments of seven clay figurines (“idols”). In addition, over 15,000 lithic artifacts were found at Brunn am Gebirge, most of them at Brunn 2, which is unusual for a central European Neolithic settlement. The presence of anthropomorphic forms, amphora, remains of musical instruments (clay flutes), as well as large quantities of lithic artifacts indicates that Brunn 2 was part of a “central settlement”, a site dedicated to the ritual activities of large LBK communities^23^.

### Brunn 2 burials

Burial 1 was found in a clay extraction pit in the southern part of the Brunn am Gebirge site 2. Burials 2 and 4 were found in the long ditches of houses no longer in use at the time of burial. Burial 3 was not associated with an above-ground structure. All four individuals were buried in a typical LBK position on their left side and had a head orientation to the northeast facing east. The teeth from all four seem to have been heavily worn, suggestive of a plant-based diet^20^. One of the skeletons (Individual 2) was found with six trapezes made from radiolarite sourced some 200 km southeast at Bakony-Szentgál, near Lake Balaton in western Hungary^19^. A detailed description of the four individuals interred at Brunn 2 can be found in^20^.

### Radiocarbon dating and stable isotope analysis

Radiocarbon measurements via Accelerated Mass Spectrometry (AMS) and carbon and nitrogen (δ^13^C and δ^15^N) stable isotope analyses were accomplished at BETA Analytic, Miami, FL (BETA) and the Penn State Radiocarbon ^14^C Laboratory, University Park, PA (PSUAMS).

### Strontium isotopes from enamel analysis

It is possible to obtain specific clues about the movement of people in the past from the chemistry of prehistoric human teeth. The basic principle for the isotopic proveniencing of human remains essentially involves the comparison of isotope ratios in human tooth enamel with local, or baseline, levels from the place of burial. Tooth enamel is a remarkable repository of childhood environment. Tooth enamel forms in the first years of life and remains unchanged through life and often for a very long period after death. A variety of studies have demonstrated that enamel is highly resistant to post-mortem contamination^24–26^. Enamel is largely a mineral, hydroxyapatite, Ca_10_(PO_4_)_6_(OH)_2_, composed primarily of calcium and oxygen. A few other elements also can be deposited in the apatite. Strontium and lead, for example, substitute for calcium during mineral formation.

We lightly abrade the surface of the enamel to be sampled using a dental drill to remove surficial dirt and calculus and the outermost enamel due to the possibility of contamination by diffusion. After abrading the surface, we remove one or more small chips from the side of the molar or drill 5 to 10 milligrams of powder from the enamel. Any remaining dentine is removed. Samples of enamel weighing 3-5 mg are dissolved in 5-molar nitric acid for strontium analysis. The strontium fraction is purified using EiChrom Sr-Spec resin and eluted with nitric acid followed by water. Isotopic compositions are obtained on the strontium fraction using a VG (Micromass) Sector 54 thermal ionization mass spectrometer (TIMS). Strontium is placed on single Re filaments and analyzed using a quintuple-collector dynamic mode of data collection. Internal precision for ^87^Sr/^86^Sr analyses is typically 0.0006 to 0.0009 percent standard error, based on 100 dynamic cycles of data collection i.e., ±0.000006. The analysis was performed at the Geo Chemistry Labs, University of North Carolina at Chapel Hill, Chapel Hill, NC.

### Ancient DNA procedures

Powder was obtained from teeth of all four individuals interred at Brunn 2 in a dedicated clean room. DNA was extracted^27,28^ using between 69 and 82 mg of powder, and a portion of the extract was converted into DNA sequencing libraries. For each of the 4 samples an individually barcoded and UDG treated library^29^ was built (L1). For one sample (Individual 1, S6912) additional libraries were prepared from the same DNA extract using 4 different protocols (S6912.E1.L2 non-UDG treated double-stranded, S6912.E1.L3 non-UDG treated single-stranded^30^, S6912.E1.L5 UDG treated double-stranded, S6912.E1.L6 UDG treated single-stranded). Libraries were then enriched for both the mitochondrial genome^31^ and about 1.2 million single nucleotide polymorphisms (SNPs)^32^, and sequenced on an Illumina NextSeq500 instrument. We followed a previously described bioinformatics procedure^9^, merging sequences overlapping by at least 15 base pairs, mapping to the mitochondrial genome reference sequence *rsrs* and to the human genome reference sequence *hg19* using *bwa* (v.0.6.1)^33^, and removing duplicated sequences that mapped to the same start and stop locations and had the same molecular barcodes. For mitochondrial genome analysis, we built a consensus sequence^34^, and for nuclear genome analysis, we represented each targeted SNP by one randomly chosen sequence passing previously reported minimum mapping and base qualities^9^. We evaluated ancient DNA authenticity by tabulating characteristic C-to-T damage rates at the terminal nucleotides of sequencing reads and by measuring apparent heterozygosity rates in haploid genome regions^35,36^. Full technical information on the data we produced for each sample is given in Supplementary Table S1 online.

### Whole-genome statistical analysis

Principal component analysis (PCA) was carried out using the smartpca software^37^. We computed axes using 1035 present-day individuals with West Eurasian ancestry genotyped at 593,124 SNPs on the Affymetrix Human Origins array^38^, and projected the newly reported and previously published ancient individuals^2,3,9–11,13,39–43^ using the least-squares option (‘lsqproject: YES’). We inferred patterns of shared ancestry with WHGs using *f*-statistics as previously described^3^. To test for symmetry of Individual #3 to different Neolithic populations, we used the statistic *f*_*4*_(#3, Mbuti; Neolithic_A, Neolithic_B), with Central African hunter-gatherers as an outgroup^44^.

## RESULTS

### Genetic analyses

We obtained genetic data passing quality control for three out of the four individuals interred at Brunn 2. No usable genetic data was obtained from Individual 4. Individuals 1-3 were males by genetic typing. The mitochondrial lineages of Individuals 1-3 were J1, U5a1, and K1b1a (Table 1), while their Y chromosomal lineages were BT, CT, and G2a2a1a, respectively. For Individual 1, we note that we ostensibly observed derived alleles at the diagnostic haplogroup P sites CTS3446 and F212, the R1 site CTS997, and the R1b1a1a2 sites PF6444 and L749 (nomenclature from the International Society of Genetic Genealogy, http://www.isogg.org), but these were mostly carried on long sequencing reads (41, 96, 74, 131, and 96 bases, respectively), none of which had evidence of ancient DNA damage, so we believe some or all of them to be due to low levels of contamination. We also observed an ancestral allele at the haplogroup R site L1225 (read length 45, likewise not damaged).

**Table 1.**
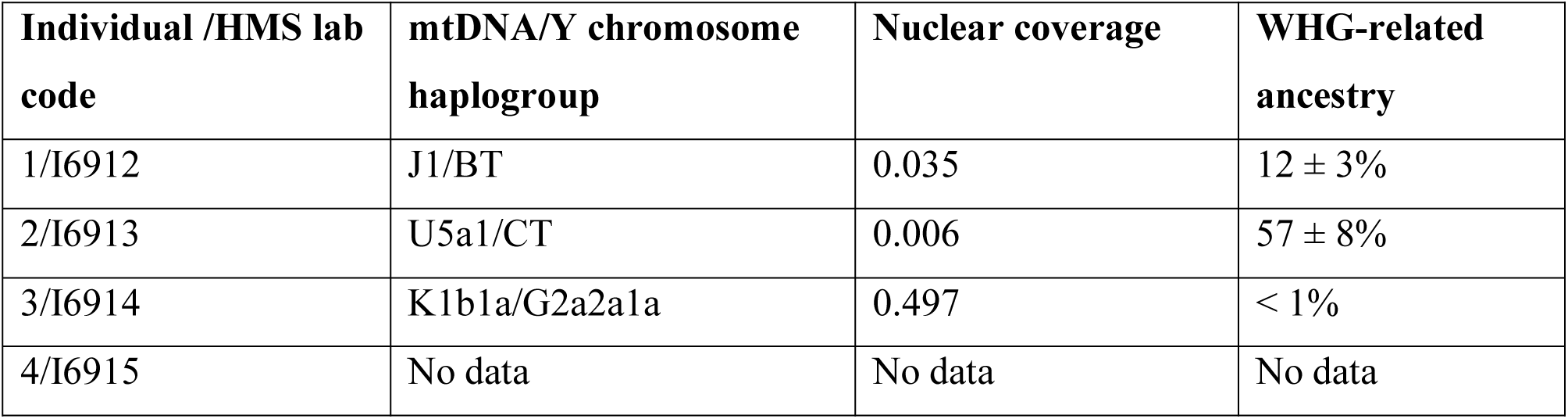
Genetic data for Brunn 2 individuals.

SNP data obtained from whole genome sequencing were used as the basis for a PCA plot comparing to Neolithic Anatolian and early Neolithic European individuals (Figure 2). Individual 2 (I6913) fell closest to WHGs but shifted toward EEFs/ANFs, while Individuals 1 (I6912) and 3 (I6914) grouped with Anatolian Neolithic farmers and closely related central European farming groups. In the zoomed-in plot (Supplementary Fig. S1 online), I6914 appears to be borderline between ANFs and ENFs, while I6912 is on the high end of WHG relatedness for early European farmers. We note though that I6912 and especially I6913 have relatively low sequencing coverage, so their exact positions in PCA should be interpreted with caution.

**Figure 2.**
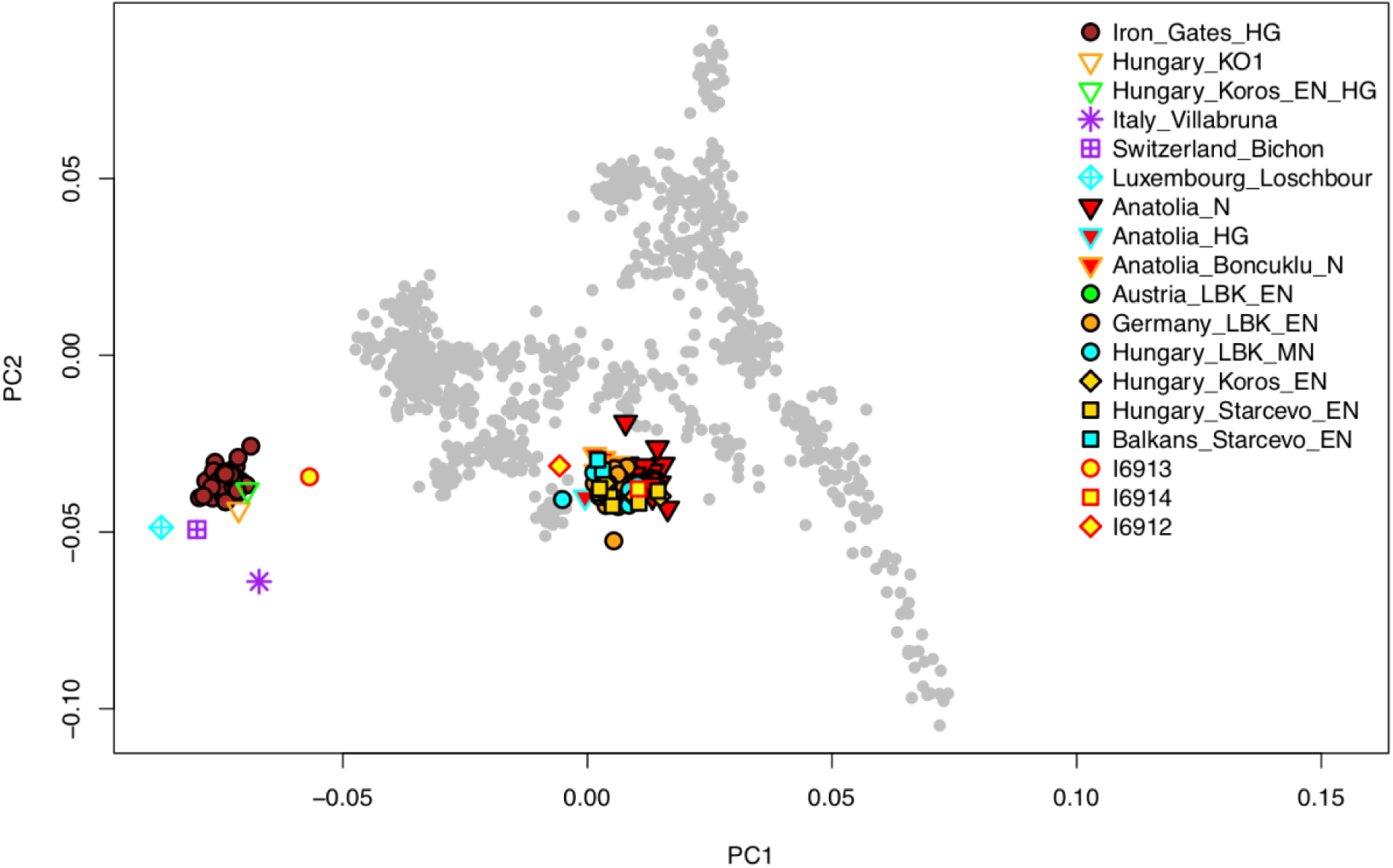
Individuals from Brunn am Gebirge site 2 plotted onto the space of principle components defined by 1035 present-day individuals with West Eurasian ancestry and including selected published ancient individuals of European and Anatolian hunter gatherer and Anatolian farming ancestries. Sources of the published data are given in the Materials and Methods. I6912, Individual 1; I6913, Individual 2; I6914, Individual 3. The PCA computation was done with smartpca, version 16690 and the visualization was made in R, version 3.5.1.

To formalize these observations, we used *f*-statistics to measure asymmetries in allele sharing (see Materials and Methods). For Individuals 1-3, we estimated genome-wide WHG-related ancestry proportions of 12±3%, 57±8%, and <1%, respectively, consistent with the PCA results (Table 1). For Individual 1, we determined his WHG ancestry to be more closely related to western and central European hunter-gatherers than to southeastern European hunter-gatherers (*f*_*4*_(#1, Anatolia N; W/C WHG, SE WHG) = −2.0*10^−3^, |Z| = 2.4; c.f. ref. 3); for Individual 2, this distinction could not be made with precision (|Z| < 1). Finally, we tested for unequal relatedness of Individual 3 (who yielded the highest sequencing coverage) to various published Neolithic farmers and found that while he is approximately symmetrically related to Neolithic Anatolians and Starčevo-associated individuals from southeastern Europe, he does share excess alleles with other LBK groups from central Europe (|Z| > 3 for Germany, |Z| > 5 for Austria). We note that this signal is not caused by WHG-related ancestry in the LBK groups, as the statistic value for WHG has the opposite sign (*f*_*4*_(#3, Mbuti; WHG, Anatolia N) << 0, Z < −17).

### Stable isotope (δ^13^C and δ^15^N) data

Stable isotope data on bone material from Individuals 1 and 2 revealed collagen depletion, therefore, the data from those two bone samples are unreliable. The results of stable isotope analysis from dentin revealed that all three individuals grew up on a diet consistent with that of a Neolithic farming community (Table 2, Figure 3). When the of δ^13^C and δ^15^N values from the Brunn 2 individuals are plotted against the average δ^13^C and δ^15^N values from European LBK sites^45^, European Inland Neolithic farming communities (EIN) and European Inland Mesolithic HGs (EIM)^46^, as well as Anatolian Neolithic farmers (AN)^9^, all three Brunn 2 individuals place within the range of Anatolian and European Neolithic farmers (Figure 3). The values for Individual 2 from dentin and bone overlap with the lower margin for δ^15^N values of EIM, while the δ^15^N values for Individuals 1 and 2 fall outside of the EIM margin and within the EIN margin (Figure 3).

**Table 2.** Radiocarbon and stable isotope data of the individuals from Brunn am Gebirge site 2, as well as mean stable isotope values (± SD) for European inland Mesolithic (EIM) and European inland Neolithic (EIN)^46^, central European LBK^45^, and Anatolian Neolithic (AN)^9^. Radiocarbon dates for Individuals 2 and 4 (ETH-14827 and ETH-11150) are from^20^.

| Code | Sample | Material | Uncalibrated date, uncalBP | Calibrated date, calBCE (2-σ) | $\delta^{13}\text{C}$ (‰) | %C | $\delta^{15}\text{N}$ (‰) | %N | C:N ratio | $^{86}\text{Sr}/^{87}\text{Sr}$ |
| --- | --- | --- | --- | --- | --- | --- | --- | --- | --- | --- |
| BETA-506840 | Grave 1 | Bone | - | - | -21.32 | 20.46 | 9.21 | 6.38 | 3.74 |  |
| BETA - 508239 | Grave 1 | Dentin | 6,510±30 | 5,534-5,380 | -20.3 | 42.09 | 9.78 | 15.5 | 3.2 | |
| ETH-14827 | Grave 2 | Bone | 6,460±70 | 5,551-5,307 | - | - | - | - | - | |
| BETA - 506842 | Grave 2 | Dentin (molar crown) | - | - | -20.16 | 41.76 | 9.21 | 15.27 | 3.2 |  |
| BETA - 508238/<br>506841 | Grave 2 | Bone | - | - | -23.23 | 44.81 | 8.75 | 9.05 | 5.8 |  |
| PSUAMS-3468 | Grave 3 | Dentin (molar) | 6,360±30 | 5,464-5,234 | -20.31 | 46.36 | 10.35 | 16.35 | 3.31 | |
| BETA -<br>506843 | Grave 3 | Bone | - | - | -20.42 | 35.56 | 10.43 | 12.22 | 3.4 |  |
| ETH-11150 | Grave 4 | Bone | 6,360±50 | 5,470-5226 | - | - | - | - | - |  |
| F10767 | Grave 1 | Enamel |  |  |  |  |  |  |  | 0.708823 |
| F10766 | Grave 2 | Enamel |  |  |  |  |  |  |  | 0.711512 |
|  | EIM |  |  |  | -20.7±1.8 |  | 12.5±2.4 |  |  |  |
|  | EIN |  |  |  | -20.1±0.6 |  | 9.6±1.1 |  |  |  |
|  | LBK |  |  |  | -20.03<br>±0.27 |  | 9.64±0.73 |  |  |  |
|  | AN |  |  |  | -19.52<br>±0.83 |  | 10.52±1.2 |  |  |  |

**Figure 3.**
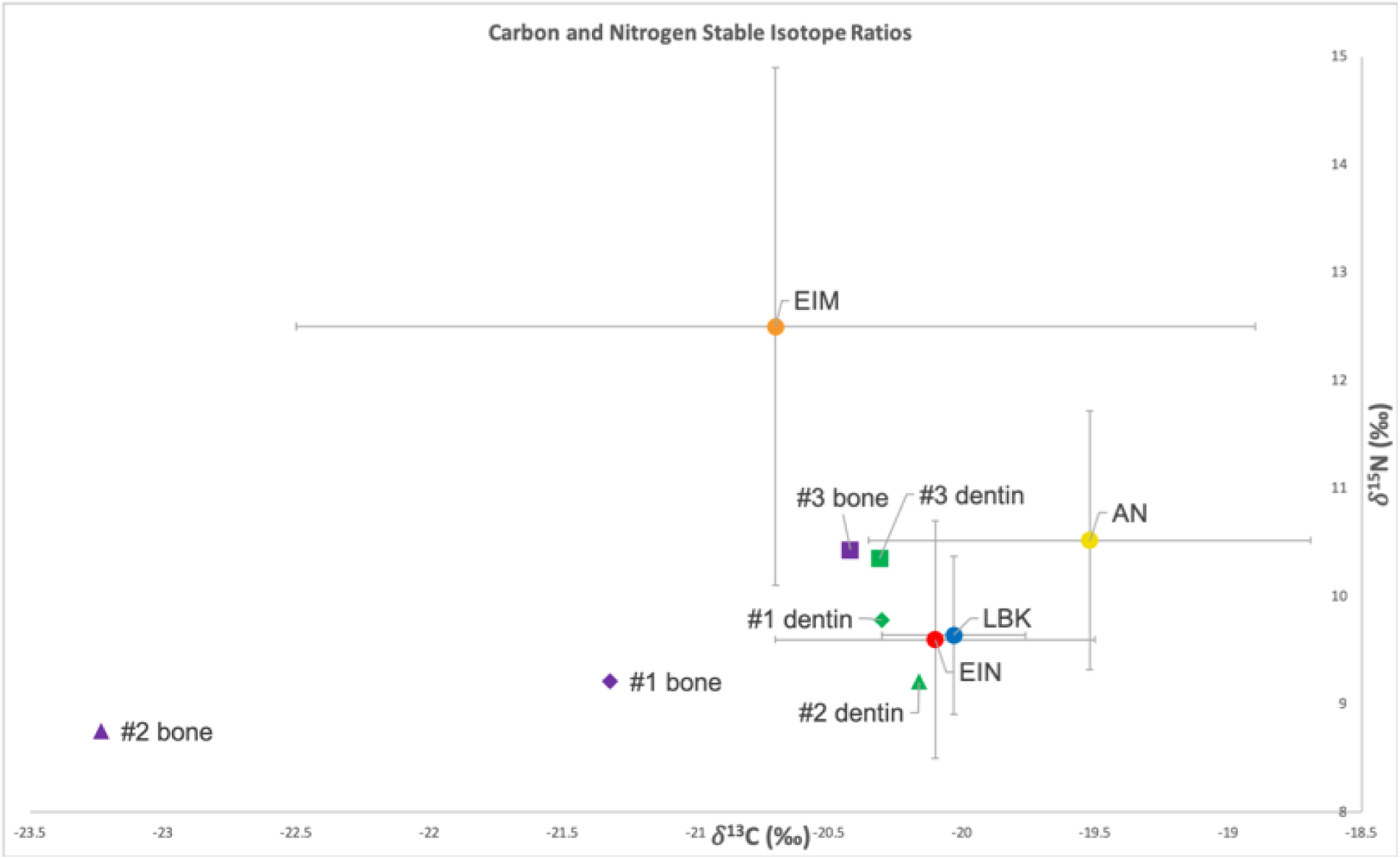
Distribution of carbon and nitrogen isotope ratios of Individuals 1, 2 and 3 from Brunn am Gebirge site 2 in relation to the mean isotope ratios (±SD) from Anatolian Neolithic (AN), European Inland Mesolithic (EIM), European Inland Neolithic (EIN), and *Linearbandkeramik* (LBK) populations. Sources of the published data are given in Table 2. The figure and underlying statistical analyses were generated using Microsoft Excel 2019 for Mac, version 16.25.

### Strontium isotope analysis

A small-scale strontium isotope analysis of two of the burials (Individuals 1 and 2) from Brunn 2 was undertaken, limited by the amount of enamel that had survived. Strontium isotope ratios (^86^Sr/^87^Sr) in human tooth enamel reflect the nutrients consumed during tissue formation in early childhood, assumed to represent the place of birth. Local or baseline ratios at the place of burial should reflect the place of death. Difference between enamel and baseline should identify individuals of non-local birth.

Baseline values were not directly available from the site of Brunn 2 but had been measured at a nearby Longobard cemetery less than 100 m from Brunn 2 (Peter Stadler, personal communication, 2019). Measurement of modern vegetation suggests a local range of ^86^Sr/^87^Sr values between 0.7085 and 0.7098.

The results of ^86^Sr/^87^Sr measurements are presented in Table 2. If we use the Longobard cemetery values in consideration of the two human burials from Brunn 2 then Individual 1 is local and Individual 2 is non-local, falling outside the local range based on the value from the nearby cemetery.

## DISCUSSION

While the remains of early Neolithic farmers in central Europe are relatively abundant, very few remains of contemporaneous hunter gatherers are known, making the understanding of the HGs’ lifeways and their integrations with incoming Anatolian-related farming migrants difficult, particularly during the earliest steps of the Neolithization of Europe. While recent genetic studies have pointed to a limited genetic exchange between immigrant farmers and local European HGs, the flow of material goods from farmers to HGs has been documented (see references in ^47^). The reciprocal material goods flow into the farming communities has been more difficult to identify^47^.

A bioarchaeological analysis of the remains of the interred at Brunn 2 presented in this study allows insight into the life history of early European Neolithic farmers who lived near the beginning of the establishment of farming economies in central Europe, revealing evidence of biological interaction between incoming Anatolians and local HGs during the earliest stages of the arrival of farming in the Neolithic Central Europe.

The mtDNA lineages of Individuals 1 and 3 belong to two of the most common mtDNA subclades found in Neolithic individuals from the Near East as well as their Neolithic European descendants^3,48,49^. At the same time, an individual belonging to K1b (K1b2) has been identified in a Mesolithic forager from the Baltic^50^. On the other hand, divisions of haplogroup U5 such as the U5a1 lineage identified in Individual 2 are generally considered to be characteristic of European hunter-gatherers^51,52^. At the same time, a U5-carrying individual (U5b2) has recently been identified in the Çatalhöyük population of central Anatolia^53^.

Individual 3 carries Y chromosome haplogroup G2a2a1a, from the larger set of G2a Y chromosomal lineages, which are characteristic of ANF and ENF populations^9,15,48^. Individuals 1 and 2 have Y-chromosomes from the macro-lineages BT and CT, respectively, but due to their low-coverage data, we are not able to assign them with greater precision.

The genetic signature of Individual 3 is that of Neolithic Anatolian-related ancestry, consistent with that of most of the representatives of European Neolithic farming cultures, including LBK and Starčevo. Our analyses indicate that this individual had very little ancestry derived from European hunter-gatherers (likely zero, and no more than 1%). At the same time, Individuals 1 and 2 had WHG ancestry that was acquired after their ancestors had left Anatolia. We were unable to determine dates for this admixture, so it is possible either that it occurred locally or that the Brunn 2 migrants encountered WHGs along their journey and integrated WHG ancestry into the their predominantly ANF-derived genetic pool prior to their arrival in central Europe. It is also possible that the Brunn 2 migrants interacted with, or descended from, the Anatolia-derived farming communities (monochrome white painted pottery groups) that settled in the Balkans ca. 600 years earlier and would have also had opportunities to incorporate WHG ancestry in their gene pool since leaving Anatolia. However, the very high WHG-related proportion in Individual 2, combined with the western European affinity of the HG-related ancestry in Individual 1, points toward recent post-arrival admixture in central Europe as the most likely scenario.

Lithic artifacts from Brunn 2 indicate active interaction between early ENF and local HG population groups in the Early Neolithic. A likely explanation of the presence of 15,000 lithic artifacts at the Brunn am Gebirge - Wolfholz site is that the Neolithic farmers produced hunting implements for trade with local HGs. The absence of the evidence of violence at early LBK settlements suggests low hostility between the local HGs and LBK farmers^47^. The material for lithic implements from the grave of Individual 2 was sourced from around Lake Balaton^20^ where other settlements of the Formative LBK phase have been found. Individual 2’s strontium isotope ratio measurements indicate he was not born at the Brunn 2 settlement site and could conceivably have come from the area where the lithic material had been procured, namely Bakony-Szentgál in Hungary. The sourcing of the lithic material from distal areas is not uncommon for LBK settlements in Austria^54^, but the number of lithic artifacts found at Brunn 2 may have significance. A lithic production center at Brunn 2 would have needed the knowledge of dedicated craftsmen, some of them could conceivably have come from the local HG communities. This might explain the presence of individuals such as Individual 2, with high proportions of HG-related ancestry, on a permanent or a semi-permanent basis within LBK settlements, and their subsequent integration into the LBK communities.

The diet isotope δ^15^N ratios from dentin ranged from 9.21 for Individual 2 to 9.78 for individual 1 to 10.35 for Individual 3. Individuals 1 and 2 are roughly contemporaneous, and their δ^15^N variation likely reflects individual dietary specifics. However, the δ^15^N value for Individual 3 is somewhat elevated compared to that of Individuals 1 and 2. The δ^15^N value for Individual 3 is in the upper range for δ^15^N variation for EIN and LBK, within the lower range of δ^15^N variation for EIM, and within the average for δ^15^N of ANF (Table 2, Figure 3). Individual 3 is also chronologically the youngest of the three. As with Individuals 1 and 2, the δ^15^N measurements in the Individual 3 could reflect the specifics of individual dietary patterns. At the same time, the varying nitrogen isotope ratios could be a result of oscillating environmental conditions during the Formative phase of LBK leading to the failure of initially maladapted domesticated plants from semi-arid Anatolia to thrive in the continental climate of central Europe^55^ and causing early European farmers to periodically rely more on animal protein rather than agricultural crops. Another explanation for the δ^15^N variation could be the progressive expansion of domestic animal herds in early LBK leading to an increased availability of animal protein and/or increased crop manuring, also leading to elevated δ^15^N values^55^. Lastly, the elevated δ^15^N values of Individual 3 could also be connected with his occupation, of which we do not have sufficient information.

The symmetric genetic relationship of Individual 3 to Starčevo and ANF individuals studied to date and his greater genetic affinity to other LBK individuals implies that the individual was from a population that had experienced a small amount of genetic drift not shared with Anatolian and southeastern European farmers studied to date. In theory, the excess relatedness of Individual 3 to other LBK-associated individuals could be due to shared WHG ancestry, but (a) we find approximately zero such ancestry in Individual 3, and (b) direct allele-sharing tests show that such a signal would in fact be in the opposite direction. In any case, our results show that the lineage that gave rise to the primary ancestry of central European LBK-associated populations was represented at Brunn 2 together with other sampled early Neolithic sites.

It is clear that the process of formation of the Starčevo and Linear Pottery cultures was more complicated than a mere immigration into a new area and the subsequent cultural deterioration during the movement. It had to also include the influence of local populations (the Early Neolithic in Bulgaria, Serbia and Croatia, and the Mesolithic in Hungary and Austria), and the adaptation to new ecological conditions, as well as new sources of stone, clay etc. We can thus conclude that the migration model of the European Neolithization involved the movement of the carriers of the agrarian economy from Anatolia, who were variably influenced by either the Mesolithic or Neolithic populations from earlier migration events already living in the Balkans, which then established the LBK culture once they arrived at Brunn 2 and other sites of the formative LBK phase. The finding of remains of a possible first generation ANF/WHG admixed individual interred at Brunn 2 points to the economic, cultural and biological integration of HGs into the early LBK farming community. The full extent of contribution of European HGs to incoming Anatolian farmers remains an important subject for future work.

## Acknowledgments

We thank Nasreen Broomandkhoshbacht, Matthew Ferry, Megan Michel, Jonas Oppenheimer, and Kristin Stewardson for help with ancient DNA laboratory work; Matthew Mah and Swapan Mallick for help with bioinformatics; and Iñigo Olalde and Vagheesh Narasimhan for help with Y chromosome analysis. D.R. is an Investigator of the Howard Hughes Medical Institute and his ancient DNA laboratory work was supported by National Science Foundation HOMINID grant BCS-1032255, by National Institutes of Health grant GM100233, by an Allen Discovery Center grant, and by grant 61220 from the John Templeton Foundation.

## References

1. Lukes, A. & Zvelebil, M. Inter-generational transmission of culture and LBK origins: Some indications from eastern-central Europe. in Living Well Together? Settlement and Materiality in the Neolithic of South-East and Central Europe (eds. Bailey, D., Whittle, A. & Hofmann, D.) 139–150 (Oxbow Books, 2008).

2. Gamba, C. et al. Genome flux and stasis in a five millennium transect of European prehistory. Nat. Commun. 5, 5257 (2014).

3. Lipson, M. et al. Parallel palaeogenomic transects reveal complex genetic history of early European farmers. Nature 551, 368 (2017).

4. Childe, V. The dawn of European civilisation. (Kegan Publisher, 1957).

5. Piggott, S. E. Ancient Europe from the Beginnings of Agriculture to Classical Antiquity: A Survey. (Edinburgh University Press, 1965).

6. Vencl, S. The role of hunting-gathering populations in the transition to farming, a Central European perspective. in Hunters in transition: Mesolithic societies of temperate Eurasia and their transition to farming (ed. Zvelebil, M.) 43–52 (Cambridge University Press, 1986),

7. Cavalli-Sforza, L. L. & Cavalli-Sforza, F.. The great human diasporas: The history of diversity and evolution. (Addison-Wesley, 1995).

8. Van Andel, T. H. & Runnels, C. N. The earliest farmers in Europe. Antiquity 69, 481–500 (1995).

9. Mathieson, I. et al. Genome-wide patterns of selection in 230 ancient Eurasians. Nature 528, 499–503 (2015).

10. Kılınç, G. M. et al. The Demographic Development of the First Farmers in Anatolia. Curr. Biol. (2016). doi:10.1016/j.cub.2016.07.057

11. Lazaridis, I. et al. Ancient human genomes suggest three ancestral populations for present-day Europeans. Nature 513, 409–413 (2014).

12. Brandt, G. et al. Ancient DNA reveals key stages in the formation of central European mitochondrial genetic diversity. Science 342, 257–61 (2013).

13. Haak, W. et al. Massive migration from the steppe was a source for Indo-European languages in Europe. Nature 522, 207–11 (2015).

14. Günther, T. et al. Ancient genomes link early farmers from Atapuerca in Spain to modern-day Basques. Proc. Natl. Acad. Sci. 112, 11917–11922 (2015).

15. Hofmanová, Z. et al. Early farmers from across Europe directly descended from Neolithic Aegeans. Proc. Natl. Acad. Sci. 113, 6886–6891 (2016).

16. Stadler, P. & Kotova, N. Early Neolithic settlement from Brunn Wolfholz in Lower Austria and the problem of the origin of (Western) LBK. in Neolithization of the Carpathian Basin: Northernmost distribution of the Starčevo-Körös culture (eds. Kozłowski, J.. & Raczky, P.) 325–348 (Polish Academy of Arts and Sciences, 2010).

17. Stadler, P. & Kotova, N. The Early LBK Site at Brunn am Gebirge, Wolfholz (5670-5100 BC); Locally Established or Founded by Immigrants from the Starčevo Territory? in Ősrégészeti Tanulmányok / Prehistoric Studies 1 (eds. Anders, A. & Kulcsar, G.) 259–275 (L’Harmattan, 2013).

18. Stadler, P. & Kotova, N. Early Neolithic Settlement from Brunn Wolfholz in Lower Austria and the problem of the Origin of the (Western) LBK. in The Domestic Space in LBK Settlements. Internationale Archäologie Arbeitsgemeinschaft 17 (eds. Hamon, C., Allard, P. & Ilett, M.) 51–78 (Verlag Marie Leidorf, 2013).

19. Stadler, P. Settlement of the Early Linear Ceramics Culture at Brunn am Gebirge, Wolfholz site. Doc. Praehist. 32, 269–278 (2005).

20. Early Neolithic Settlement Brunn am Gebirge, Wolfholz, in Lower Austria. Volume I. Early Neolithic Settlement Brunn am Gebirge, Wolfholz, Site 2 in Lower Austria and the Origin of the Western Linear Pottery Culture (LPC). (Beier & Beran. Archaologische Fachliteratur, 2019).

21. Bánffy, E. The 6th millennium BC boundary in Western Transdanubia and its role in the Central European Transition. The Szentgyörgyvölgyi-Pityerdomb settlement. (Archaeological Institute of the HAS, 2004).

22. Simon, K. Das Fundmaterial der frühesten Phase der transdanubischen Linienbandkeramik auf dem Fundort Zalaegerszeg-Andráshida// Gébárti-tó// Arbeitsplatz III. Antaeus 025, 189–203 (2002).

23. Kotova, N. & Stadler, P. Sites in Brunn-am-Gebirge and the Problem of the Hierarchy of the Oldest Linear Pottery Culture Settlements. Strat. Plus 2, 109–117 (2018).

24. Budd, P., Montgomery, J., Barreiro, B. & Thomas, R. G. Differential diagenesis of strontium in archaeological human dental tissues. Appl. Geochemistry (2000). doi:10.1016/S0883-2927(99)00069-4

25. Kohn, M. J., Schoeninger, M. J. & Barker, W. W. Altered states: Effects of diagenesis on fossil tooth chemistry. Geochim. Cosmochim. Acta (1999). doi:10.1016/S0016-7037(99)00208-2

26. Lee-Thorp, J. & Sponheimer, M. Three case studies used to reassess the reliability of fossil bone and enamel isotope signals for paleodietary studies. J. Anthropol. Archaeol. (2003). doi: 10.1016/S0278-4165(03)00035-7

27. Dabney, J. et al. Complete mitochondrial genome sequence of a Middle Pleistocene cave bear reconstructed from ultrashort DNA fragments. Proc. Natl. Acad. Sci. 110, 15758–15763 (2013).

28. Korlević, P. et al. Reducing microbial and human contamination in DNA extractions from ancient bones and teeth. Biotechniques 59, (2015).

29. Rohland, N., Harney, E., Mallick, S., Nordenfelt, S. & Reich, D. Partial uracil –DNA – glycosylase treatment for screening of ancient DNA. Philos. Trans. R. Soc. B Biol. Sci. (2015). doi:10.1098/rstb.2013.0624

30. Gansauge, M.-T. et al. Single-stranded DNA library preparation from highly degraded DNA using T4 DNA ligase. Nucleic Acids Res. gkx033 (2017). doi:10.1093/nar/gkx033

31. Maricic, T., Whitten, M. & Pääbo, S. Multiplexed DNA Sequence Capture of Mitochondrial Genomes Using PCR Products. PLoS One 5, e14004 (2010).

32. Fu, Q. et al. An early modern human from Romania with a recent Neanderthal ancestor. Nature 524, 216–219 (2015).

33. Li, H. & Durbin, R. Fast and accurate long-read alignment with Burrows-Wheeler transform. Bioinformatics (2010). doi:10.1093/bioinformatics/btp698

34. Weissensteiner, H. et al. HaploGrep 2: mitochondrial haplogroup classification in the era of high-throughput sequencing. Nucleic Acids Res. (2016). doi:10.1093/nar/gkw233

35. Fu, Q. et al. DNA analysis of an early modern human from Tianyuan Cave, China. Proc. Natl. Acad. Sci. 110, 2223–2227 (2013).

36. Korneliussen, T. S., Albrechtsen, A. & Nielsen, R. ANGSD: Analysis of Next Generation Sequencing Data. BMC Bioinformatics 15, 356 (2014).

37. Patterson, N., Price, A. L. & Reich, D. Population structure and eigenanalysis. PLoS Genet. (2006). doi:10.1371/journal.pgen.0020190

38. Lazaridis, I. et al. Genetic origins of the Minoans and Mycenaeans. Nature (2017). doi:10.1038/nature23310

39. Feldman, M. et al. Late Pleistocene human genome suggests a local origin for the first farmers of central Anatolia. Nat. Commun. 10, 1218 (2019).

40. Olalde, I. et al. Derived immune and ancestral pigmentation alleles in a 7,000-year-old Mesolithic European. Nature 507, 225 (2014).

41. Jones, E. R. et al. Upper Palaeolithic genomes reveal deep roots of modern Eurasians. Nat. Commun. 6, 8912 (2015).

42. Fu, Q. et al. The genetic history of Ice Age Europe. Nature 534, 200–205 (2016).

43. Mathieson, I. et al. The genomic history of southeastern Europe. Nature 555, 197–203 (2018).

44. Mallick, S. et al. The Simons Genome Diversity Project: 300 genomes from 142 diverse populations. Nature (2016). doi:10.1038/nature18964

45. Ash, A. et al. Regional differences in health, diet and weaning patterns amongst the first Neolithic farmers of central Europe. Sci. Rep. 6, 29458 (2016).

46. Schulting, R. Dietary Shifts at the Mesolithic–Neolithic Transition in Europe: An Overview of the Stable Isotope Data. in The Oxford Handbook of the Archaeology of Diet (eds. Lee-Thorp, J. & Katzenberg, M. A.) 1–32 (Oxford University Press, 2018). doi:10.1093/oxfordhb/9780199694013.013.35

47. Vanmontfort, B. Forager–farmer connections in an ‘unoccupied’ land: First contact on the western edge of LBK territory. J. Anthropol. Archaeol. 27, 149–160 (2008).

48. Lazaridis, I. et al. Genomic insights into the origin of farming in the ancient Near East. Nature 536, 419–424 (2016).

49. Szécsényi-Nagy, A. et al. Tracing the genetic origin of Europe’s first farmers reveals insights into their social organization. Proc. Biol. Sci. 282, 20150339 (2015).

50. Mittnik, A. et al. The genetic prehistory of the Baltic Sea region. Nat. Commun. 9, 442 (2018).

51. Bramanti, B. et al. Genetic Discontinuity Between Local Hunter-Gatherers and Central Europe’s First Farmers. Science (80-.). 326, 137–140 (2009).

52. Malmström, H. et al. Ancient mitochondrial DNA from the northern fringe of the Neolithic farming expansion in Europe sheds light on the dispersion process. Philos. Trans. R. Soc. Lond. B. Biol. Sci. 370, 20130373 (2015).

53. Chyleński, M. et al. Ancient Mitochondrial Genomes Reveal the Absence of Maternal Kinship in the Burials of Çatalhöyük People and Their Genetic Affinities. Genes (Basel). 10, 207 (2019).

54. Neugebauer-Maresch, C. & Lenneis, E. Origin and contacts of people buried at the LBK graveyard at Kleinhadersdorf, Austria. Doc. Praehist. 40, 305–312 (2013).

55. Ivanova, M., De Cupere, B., Ethier, J. & Marinova, E. Pioneer farming in southeast Europe during the early sixth millennium BC: Climate-related adaptations in the exploitation of plants and animals. PLoS One 13, e0197225 (2018).

