## Supplementary Figure 1 for "Interactions between earliest *Linearbandkeramik* farmers and central European hunter gatherers at the dawn of European Neolithization"

Supplementary Figure S1. A zoomed-in PCA plot from Figure 1, showing the position of Individuals 1 (I6912) and 3 (I6914) relative to Anatolian and European Neolithic farmers.

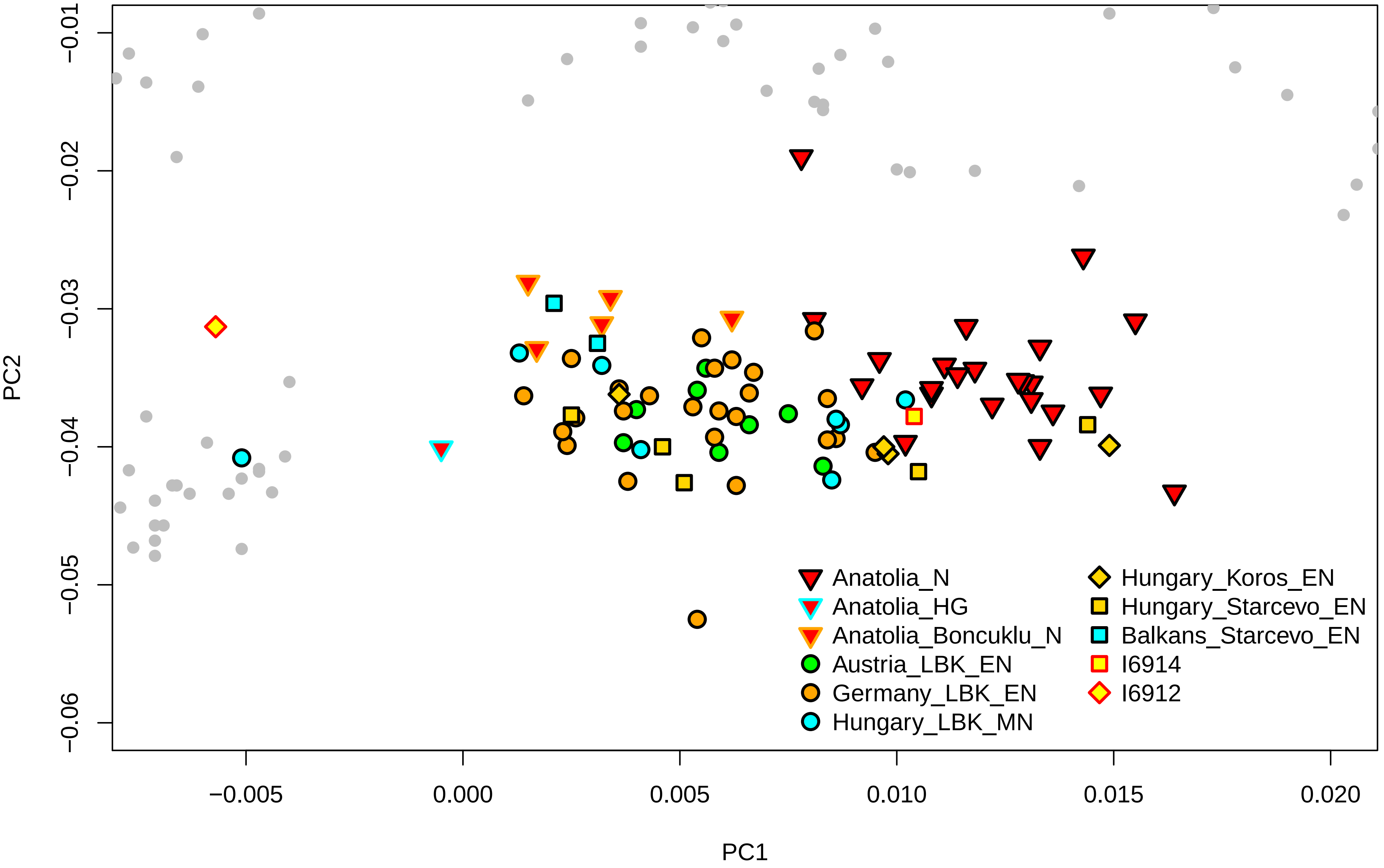
