## Supplementary Table 1 for "Interactions between earliest *Linearbandkeramik* farmers and central European hunter gatherers at the dawn of European Neolithization"

| Library ID | Lab Ind. ID | Skeletal Code/Element | Date | Locality | Country | Latitude | Longitude | mg Powder Extracted | Library protocol | UDG Treatment | uL Extract Used | mg Equivalent Powder Used | mtDNA Coverage |
|---|---|---|---|---|---|---|---|---|---|---|---|---|---|
| S6912.E1.L1 | I6912 | Ind. #1, left lower first or second molar ( | 5515-5370 calBCE (6510±30 BP, Beta-5( | Brunn Wolfholz | Austria | 48.12 | 16.29 | 69 | Double-stranded | partial | 10 | 7.7 | 54.4 |
| S6912.E1.L2 | I6912 | Ind. #1, left lower first or second molar ( | 5515-5370 calBCE (6510±30 BP, Beta-5( | Brunn Wolfholz | Austria | 48.12 | 16.29 | 69 | Double-stranded | minus | 10 | 7.7 | 55.5 |
| S6912.E1.L3 | I6912 | Ind. #1, left lower first or second molar ( | 5515-5370 calBCE (6510±30 BP, Beta-5( | Brunn Wolfholz | Austria | 48.12 | 16.29 | 69 | Single-stranded | minus | 10 | 7.7 | 127.3 |
| S6912.E1.L5 | I6912 | Ind. #1, left lower first or second molar ( | 5515-5370 calBCE (6510±30 BP, Beta-5( | Brunn Wolfholz | Austria | 48.12 | 16.29 | 69 | Double-stranded | partial | 10 | 7.7 | 39.7 |
| S6912.E1.L6 | I6912 | Ind. #1, left lower first or second molar ( | 5515-5370 calBCE (6510±30 BP, Beta-5( | Brunn Wolfholz | Austria | 48.12 | 16.29 | 69 | Single-stranded | partial | 10 | 7.7 | 111.8 |
| S6913.E1.L1 | I6913 | Ind. #2, left or right lower second molar | 5551-5307 calBCE (6460±70 BP, ETH-14 | Brunn Wolfholz | Austria | 48.12 | 16.29 | 73 | Double-stranded | partial | 10 | 8.1 | 187.0 |
| S6914.E1.L1 | I6914 | Ind. #3, right lower second molar (perma | 5464-5234 calBCE (6360±30 BP, PSUAM | Brunn Wolfholz | Austria | 48.12 | 16.29 | 82 | Double-stranded | partial | 10 | 9.1 | 48.7 |
| S6915.E1.L1 | I6915 | Ind. #4, left lower second molar (perman | 5470-5226 calBCE (6360±50 BP, ETH-11 | Brunn Wolfholz | Austria | 48.12 | 16.29 | 76 | Double-stranded | partial | 10 | 8.4 | 0.9 |

| Library ID | mtDNA Median [Mean] Seq Length | mtDNA Damage Last Base | mtDNA Consensus Match | mtDNA Consensus Match 95CI | mtDNA Haplogroup | Nuclear Unique SNPS Hit | Nuclear Coverage at Targeted Positions | Nuclear Median [Mean] Seq Length | Nuclear Damage Last Base | Nuclear X Hits | Nuclear Y Hits | Nuclear Sex | Nuclear ANGSD SNPs | Nuclear ANGSD Mean | Nuclear ANGSD Z-score |
|---|---|---|---|---|---|---|---|---|---|---|---|---|---|---|---|
| S6912.E1.L1 | 45 | 0.13 | 1.000 | [0.997,1.000] | J1 | 4830 | 0.004 | 43 | 0.03 | 103 | 73 | M | 1 | | |
| S6912.E1.L2 | [53.6] | 0.45 | 0.984 | [0.963, 0.994] | J1 | 3866 | 0.004 | [58.8] | 0.32 | 77 | 40 | M | 0 | | |
| S6912.E1.L3 | [57.1] | 0.47 | 0.955 | [0.932, 0.973] | J1c8 | 10888 | 0.011 | [64.7] | 0.32 | 303 | 142 | M | 0 | | |
| S6912.E1.L5 | [50.1] | 0.13 | 0.998 | [0.990, 1.000] | J1 | 3351 | 0.003 | [55.9] | 0.09 | 56 | 25 | M | 1 | | |
| S6912.E1.L6 | [52.0] | 0.14 | 0.976 | [0.962, 0.986] | J1c | 15488 | 0.013 | [62.4] | 0.08 | 333 | 135 | M | 4 | | |
| S6913.E1.L1 | 50 | 0.08 | 1.000 | [0.998,1.000] | U5a1 | 7223 | 0.006 | 39 | 0.03 | 150 | 107 | M | 0 | | |
| S6914.E1.L1 | 43 | 0.14 | 0.961 | [0.951,0.968] | K1b1a | 429000 | 0.497 | 44 | 0.10 | 9551 | 7109 | M | 345 | 0.0044 | 0.8999 |
| S6915.E1.L1 | 43 | 0.11 | 0.918 | [0.838,0.967] | .. | .. | .. | 37 | 0.01 | .. | .. | .. | 0 | | |

**Supplementary Table S1. Library-level metrics for Brunn 2 genetic data.**
